# Comprehensive analysis using DNA metabarcoding, PCR, and HPLC unveils the adulteration in Brahmi herbal products

**DOI:** 10.1101/2022.07.30.501660

**Authors:** Abhi P. Shah, Tasnim Travadi, Sonal Sharma, Ramesh Pandit, Chaitanya Joshi, Madhvi Joshi

## Abstract

**Background:** The herbal products market is expanding and creating a bottleneck for raw materials. Hence, economically motivated adulteration has a high prevalence. DNA barcoding and species-specific PCR assays are now revolutionising the molecular identification of herbal products and are included in a number of pharmacopoeias for the identification of raw materials. High-throughput sequencing with barcoding advances toward metabarcoding, which enables the identification of unintentionally or intentionally unlabelled plant material present in herbal products. Brahmi is one of the most commercially significant and nootropic botanicals, with great controversy over the terms “Brahmi” being used to describe both *Bacopa monneri* (BM) and *Centella asiatica* (CA) species.

**Purpose:** This study evaluates DNA-based methods for Brahmi herbal products with the traditional HPLC-based analytical approach in order to assess their effectiveness.

**Methods:** We employed a species-specific PCR assay, DNA metabarcoding using *rbcL* minibarcode, and HPLC to detect the presence of the Brahmi (either BM or CA) in eighteen market samples. All the methods have been validated using in-house blended formulations.

**Results:** Comprehensive analysis of all three methods revealed the presence of 22.2%, 55.6%, and 50.0% of Brahmi by PCR assay, DNA metabarcoding, and HPLC, respectively, in Brahmi market formulations, whereas blended formulations only exhibited targeted plant species with all three methods.

**Conclusion:** Species-specific PCR can be used as a cost-effective and rapid method to detect the presence of the Brahmi, while in high-throughput methods, DNA metabarcoding can be used to detect the presence of widespread adulterated botanicals, and further, bioactive compounds could be detected by HPLC. These results emphasise the need for quality control of the marketed Brahmi herbal products as well as the implementation of all methodologies in accordance with fit for purpose.

## Introduction

Traditional medical knowledge was gleaned and updated through centuries of empirical testing. The rapid expansion of the traditional medicine market is a burning issue on a global scale, as the quality of botanicals is being compromised either unintentionally by substitute/adulteration due to a lack of taxonomic knowledge, different vernacular nomenclature, and cryptic taxa morphology or intentionally by economically motivated adulteration (Ichim, 2019; Raclariu et al., 2018a). The use of taxonomic and analytical techniques for the authentication of medicinal plants are widely acknowledged around the world, but the regulatory framework and guidelines for quality assessment may vary based on whether they are used as herbal products or dietary supplements (Joshi et al., 2017; Liu et al., 2018; Sahoo and Manchikanti, 2013). The resolution power of the analytical methods is greatly influenced by environmental factors, differences in processing and storage conditions. In addition to that, analytical methods cannot differentiate between an untargeted adulteration and substitution, however, the ability to accurately identify the therapeutic component makes them important for quality assurance and consumer safety (Raclariu et al., 2017). There is a lack of centralized authentication systems that provide taxon identification with high-resolution power. The predominance of species-specific PCR assays (Noh et al., 2021; Sharma and Shrivastava, 2016) DNA barcoding(Bansal et al., 2018; Seethapathy et al., 2015; Thakur et al., 2019; Vassou et al., 2016), and metabarcoding (Raclariu et al., 2017; Raclariu et al., 2017; Raclariu et al., 2018b; Seethapathy et al., 2019) based molecular authentication of medicinal plants has recently undergone a revolution, mostly because of DNA’s accessibility, consistency, and independence from tissue characteristics, age, harvesting techniques, and storage conditions. Chinese Pharmacopoeia, United States Pharmacopeia, British Pharmacopoeia, Japanese Pharmacopoeia, and Hong Kong Chinese Materia Medica are advocating DNA-based authentication around the globe (Wu and Shaw, 2022). Species-specific PCR assays and DNA barcoding can provide identification of single or targeted plants at a lower cost, while high-throughput sequencing-based metabarcoding can concurrently identify multiple taxon from the mixture of DNA obtained from herbal products.

Brahmi is herbal therapeutics mentioned in traditional medicine for the treatment of neurological and psychiatric disorders like loss of memory, cognitive deficits, and impaired mental function (Shinomol et al., 2011). Brahmi is a classic illustration of controversial drugs/sandigdha dravya as *Bacopa monnieri* L. (BM) (also known as waterhyssop, herb of grace, Brahmi, thyme-leaved gratiola, Indian pennywort) and *Centella asiatica* L. (CA) (also known as Mandukparni, Asiatic pennywort, Jalbrahmi, and Gotu kola), because of parallel evolving knowledge systems, currently used polynomial nomenclature in Sanskrit, varying perceptions in various communities, and vernacular equivalent (Keshari, 2021). The therapeutic effectiveness of both is very similar despite having different family classifications and taxonomy is due to having similar functional groups in secondary metabolites (Kashmira J Gohil, 2020). Around the globe $320 million market value estimated for Brahmi by 2026. The Asia-Pacific area is the largest trading sector for Brahmi, with India and China being major exporters (https://www.industryarc.com/Research/Brahmi-Market-Research-507354). Traditional markets, as well as Brahmi consumption, are prosperous in Northern America and Europe due to natural thus fewer side effects ideology, traditional value, and accessible availability (https://articlewire24.wordpress.com/2017/12/28/brahmi-market-size-and-industry-forecast-2025-market-shares-and-strategies-of-key-players/).

In our previous studies, we have used plant DNA metabarcoding (Pandit et al., 2021) and species-specific primers (Travadi et al., 2022) for the detection of adulteration in herbal formulations. In this study, we investigated adulteration in Brahmi market samples using three different approaches including species-specific PCR assay, metabarcoding and HPLC. The first objective of this study was to develop a rapid, affordable and simple PCR-based assay for the detection of BM or CA in Brahmi herbal products. The second objective was to detect a wide range of botanical adulterations in Brahmi herbal products using a DNA metabarcoding approach using an in-house developed rbcL minibarcode. The third objective is to employ established analytical techniques to confirm the presence of bioactive components. All of the three techniques have been validated by mimicking possible blended formulations. The aim of this study is to evaluate DNA-based methods for Brahmi herbal products by comparing them with the conventional HPLC-based analytical strategy advocated by the Indian Pharmacopoeia.

## Materials and Methods

### Sample description

*Centella asiatica* (CA) and *Bacopa monnieri* (BM) plants were collected from the Maharaja Sayajirao University (MSU), Vadodara, India with the help of taxonomists. Herbarium vouchers of both plants were developed and submitted at the institutional herbarium with the following voucher specimen IDs: CA: BG-181130-0005; BM: BG-181130-0018. Further, a total of 18 market samples of different companies tagged as ‘Brahmi churna/powder and vati’ were procured from the local market and e-commerce (Table S1).

### Preparation of simulated plant mixtures and DNA extraction

BM and CA whole plant species were shed-dried, grinded into a fine powder and mixed in proportions (w/w%) as follows. 1) BS1: 100% BM powder 2) BS2: 100% CA powder 3) BS3: 75% BM powder in 25% CA powder 4) BS4: 50% BM powder in 50% CA powder 5) BS5: 25% BM powder in 75% CA powder for preparing 100 mg of simulated blended formulations for the validation of PCR assay, metabarcoding and HPLC. Qiagen-DNeasy plant power pro kit (Qiagen, Germany) was used for extracting DNA from the BM and CA blended whole plant materials as well as from the market samples. DNA was extracted as per the manufacturer’s instruction from 100 mg BM and CA blended materials and 100 mg powder of market samples. During DNA extraction, proper precautions were taken to avoid cross-contamination. The *rbcL* gene was amplified to ensure the DNA quality using primers and thermal cycler conditions as described by Travadi et al. (2022).

### Development of PCR assay for detection of BM and CA

#### Designing of primers

For designing CA-specific primers, ITS sequences of BM and CA were retrieved from the NCBI and aligned using ClustalW. Alignment was done to find out nucleotides that differed between the two species. CA-specific primers were designed by considering polymorphic sites. Two online software, Oligocalc (Kibbe, 2007) and Primer 3 version 4.0.0 (Rozen and Skaletsky, 2000) were used to examine primer sequences for their optimal characteristics such as length, and melting temperature compatibility, GC content, hairpin formation, and secondary structure formation. The NCBI primer BLAST tool was used to examine the specificity and selectivity of the primers. For determining BM, sequence characterized amplified region (SCAR) primers designed by (Yadav et al., 2012) were used in this study.

#### Optimization of PCR assay

The annealing temperature for BM and CA primers were optimized in such a way that provided higher sensitivity with optimum specificity. A total of 20 µL reaction mixture was prepared comprising of 10 µL KAPA HiFi HotStart ReadyMix (Roche); 1 µL forward and reverse primers (5 pmol each); 2 µL total genomic DNA (10-15 ng/µL); 2 µL bovine serum albumin (BSA) (2 mg/L); nuclease-free water for the make-up final volume. A thermal cycler was run with the following conditions to optimize annealing temperature; initial denaturation at 95°C for 3 minutes, followed by 30 cycles of denaturation at 98 °C for 20 seconds, annealing temperature from 64 to 72 °C (with an interval of 2 °C) for 15 seconds and extension at 72 °C for 20 seconds, and final extension 72 °C for 20 seconds. For determining primer’s sensitivity, 0.001, 0.01, 0.1, 0.25, 0.5, and 1 ng of genomic DNA of BM and CA were amplified with optimized PCR conditions. To ensure the specificity and efficacy of optimized PCR assays, inter-species amplification using a mixer of BM and CA plant material was also executed. PCR assays were performed with BM and CA-specific primers and optimized PCR conditions using 10-15 ng of DNA from prepared simulated blended formulations. Optimized PCR conditions were further used for the detection of BM and CA in market samples.

#### DNA metabarcoding

For metabarcoding, we have designed *in-house* plant mini-barcodes targeting the *rbcL* gene and it was validated by performing Next Generation Sequencing (NGS) assay on authenticated plant mixture (data not shown here). The same primers were used in this study.

##### 2.3.1 DNA metabarcoding of blended formulations and market samples

The optimized PCR reaction setup and thermal cycling condition were used as mentioned above for metabarcoding of blended formulations and market samples. Amplified PCR products were purified using AMPure XP beads (Beckman Coulter). Quantification of purified PCR products was determined by Qubit 4.0 Fluorometer using the 1X dsDNA HS Assay Kit (Thermo Fisher Scientific, MA, USA). Purified PCR products were mixed into equimolar concentrations (100 pmol) and subjected to emulsion PCR (emPCR) for clonal amplification. Emulsion PCR was carried out using Ion 520™ & Ion 530™ Kit-OT2 reagent solutions (Thermo Fisher Scientific, MA, USA) with 400-bp chemistry as per manufacturer guideline. Enrichment of template-positive Ion Sphere™ Particles (ISPs) was carried out using the Ion OneTouch™ ES Instrument (Thermo Fisher Scientific, MA, USA). Sequencing was performed on the Ion GeneStudio™ S5 Plus (Thermo Fisher Scientific, MA, USA) using the Ion S5™ sequencing solutions and reagents and loaded on the Ion 530™ sequencing chip (Thermo Fisher Scientific). The efficacy of the metabarcoding assay was confirmed using simulated plant mixtures prepared as mentioned above.

#### DNA metabarcoding data analysis

The generated reads were filtered using PRINSEQ v0.20.4.31(Schmieder and Edwards, 2011) on the basis of average quality score and length. Reads with an average quality score Q < 20 and reads with length <300 bp or >350 bp were discarded. Obtained filtered reads of each sample were clustered using CD-HIT (Huang et al., 2010) at 99% identity. A representative sequence of each cluster having a minimum of five reads was further analyzed using NCBI-BLASTn (Altschul et al., 1990). To normalize data, the percentage of analysed reads is considered as 100% for determination of percent distribution of each plant.

### HPLC analysis

Authentic plant samples, simulated blended formulations and market samples were extracted using methanol (1:10 w/v) to determine the presence of bacoside A (chemical marker used to detect BM) and asiaticoside (chemical marker used to detect CA). Each extract was filtered through a 0.22 µm pore size syringe filter (HiMedia Laboratories, Mumbai, India) and 20 µL of the filtered extract was injected into a high-performance liquid chromatographic (HPLC). HPLC analysis was performed on the Shimadzu (Kyoto, Japan) system consisting of an LC-20AP pump with SPD-M20A photodiode array detector (PAD) and C18 analytical column (250 × 4.6 mm; i.d.: 5 μm). Analytical separations were carried out using a gradient of acetonitrile (A) and water containing 0.05% (v/v) orthophosphoric acid (B) as the mobile phase. The elution program was 0–25 min from 30:70 (A: B) to 40:60 (A: B), and 25–35 min from 40:60 (A: B) to 60:40 (A: B). The flow rate was 1.5 mL/min and detection was performed at 205 nm (Indian pharmacopoeia 2007, volume 3; Deepak et al. 2005). Commercially available reference standard bacoside A and asiaticoside (Sigma-Aldrich) was injected (20 µL) in HPLC in (1000 µg/mL) to obtain the chromatograms of both standards.

## Results and Discussion

### DNA extraction and rbcL gene amplification

The total DNA concentration ranges from 4.7–6.7 µg and 1.22-29.2 µg from the blended mixture and market samples, respectively (Table S2). Herbal products are processed and enriched with secondary metabolites such as polysaccharides, flavonoids, and polyphenols that are reportedly co-precipitated along with the sheared DNA and inhibit PCR amplification (Fazekas et al., 2009; Mishra et al., 2016a; Parveen et al., 2016). DNA integrity and quality were checked using the *rbcL* gene from all the extracted DNA. In all the samples, we were successfully able to amplify the *rbcL* gene, which revealed that the quality of extracted DNA is suitable for PCR amplification (Fig. S1).

### PCR based authentication of Brahmi market samples

The emergence and development of DNA-based technology has enabled the cost-effective, rapid, and yet sensitive and species-specific PCR assays to authenticate medicinal plants. Species-specific sequence-characterized amplified region (SCAR) markers, nuclear ribosomal internal transcribed spacer (nrITS) sequences, and different plastid markers (e.g., rbcL, trnH-psbA, and matK) have been extensively used to authenticate plant species(de Boer et al., 2015; Mishra et al., 2016b; Sharma and Shrivastava, 2016; Xin et al., 2018). Earlier studies have revealed a PCR-based assay for the authentication of herbal products. For instance, Noh et al. (2021) developed a PCR assay for the authentication of medicinal mistletoe species; Zhang et al. (2018) developed a PCR-based assay kit for the authentication of *Zaocys dhumnades* in traditional Chinese medicine. Similarly, as mentioned earlier, we have also developed a PCR assay for the detection of *Ocimum* sp. in Tulsi churna (Travadi et al. 2022).

To check the presence of either BM or CA in Brahmi products, we developed a species-specific primer for CA using ITS region (sequence of primers not shown here), and for BM, we used a SCAR-BM specific marker designed by Yadav et al. (2012). PCR assay optimization revealed that 65 °C is the optimum annealing temperature for both primer sets using KAPA HiFi HotStart ready mix (Fig. S2). The results of the sensitivity experiment demonstrated that the target DNA sequence was amplified successfully using BM and CA primers from 0.1 ng of total DNA (Fig. S3). No cross-amplification was observed when BM and CA primers were used with the blended formulation, and on the agarose gel, there was a band intensity gradient proportional to the percentage of plant present in the blended formulations (Fig. 1a; 1b). This established and validated the applicability of both primers. In addition, a PCR assay of Brahmi market samples revealed that out of 18 samples, four samples (i.e. 22.2% samples) showed amplification for the targeted species. Only two market samples with the labels 100 and 159 were found to detect CA, whereas two samples with the labels 220 and 223 had the presence of BM (Fig. 2). The absence of the target bands in the rest of the market samples could be attributed to the presence of counterfeit plant materials in substantial amounts.

**Figure 1.**
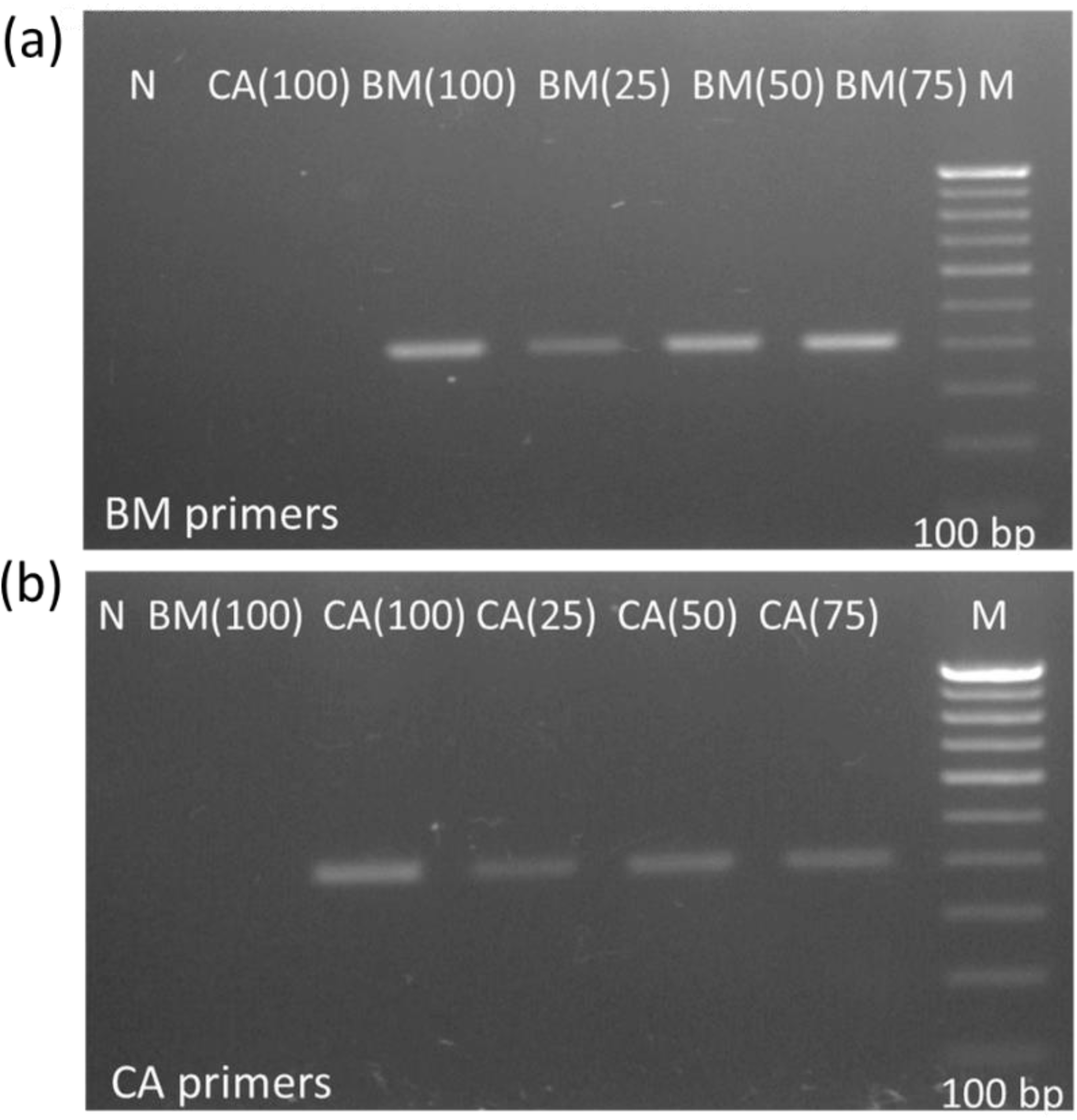
PCR assay of simulated blended formulations using *Bacopa monnieri* (BM) and *Centella asiatica* (CA) specific primers. BM (100) =100% *Bacopa monnieri*; CA (100) = 100% *Centella asiatica*; BM (75) & CA (25) = mixture of 75% of BM and 25% of CA; BM (50) & CA (50) = mixture of 50% of BM and 50% of CA; BM (25) & CA (75) = mixture of 25% of BM and 75% of CA; N= No template control; M= 100 bp ladder. (a) PCR assay of simulated blended formulations using BM primers (b) PCR assay of simulated blended formulations using CA primers

**Figure 2.**
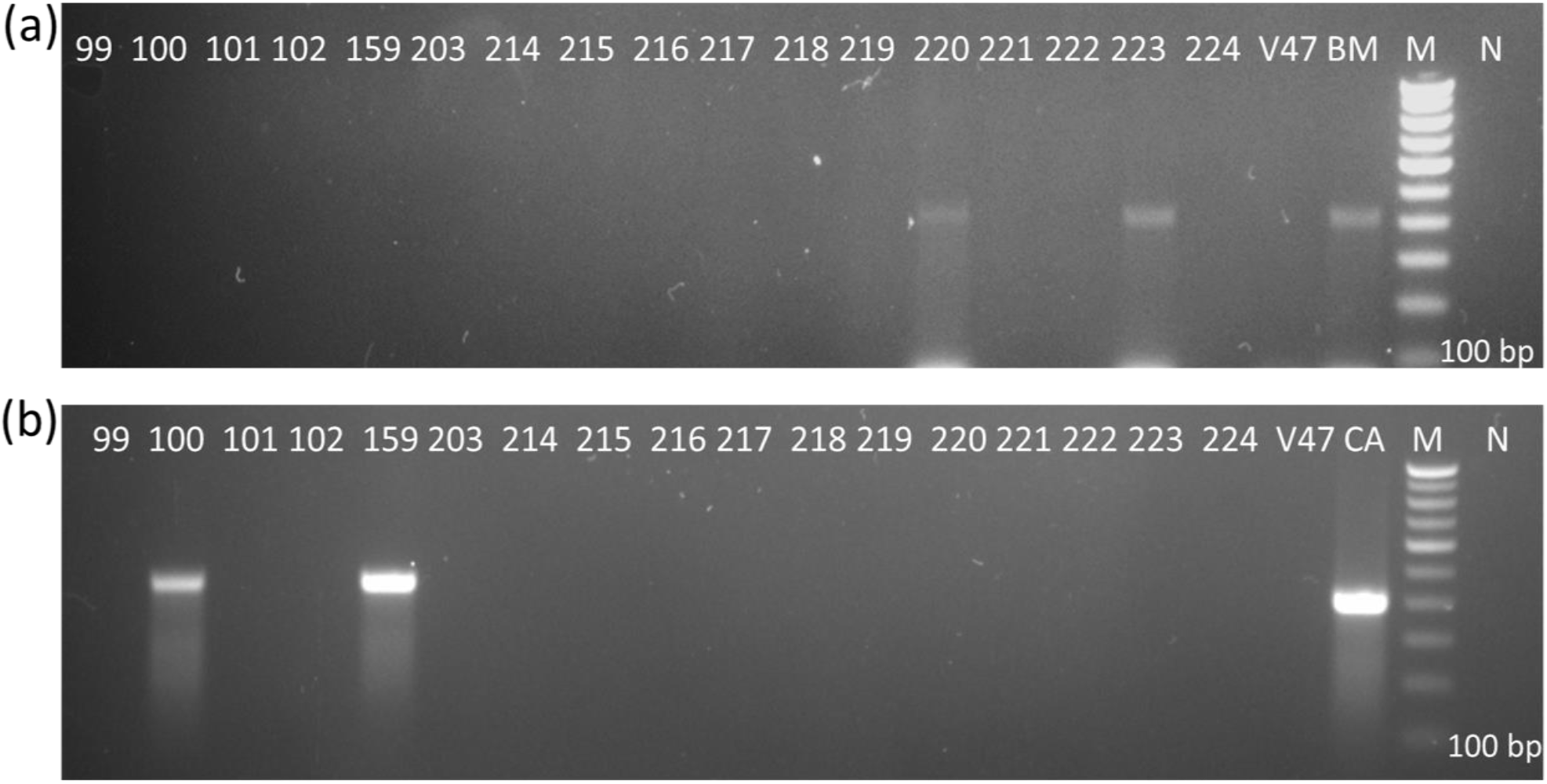
PCR assay of Brahmi market samples using *Bacopa monnieri* (BM) and *Centella asiatica* (CA) specific primers. 99, 100, 101, 102, 159, 203, 214 to 224, V47 = market sample IDs; N= No template control; M= 100 bp ladder.

### Metabarcoding of simulated blended formulations and market samples

The raw data consists of a total of 138127 reads with a mean read length of 311 bp. A total of 105353 (76.27%) reads passed our filtering quality criteria, and the mean read length of filtered reads was 338 bp (Table 1). Table 1 shows the total raw reads and percentage of reads acquired after filtering with the mean read length of each sample, as well as the percentage of analysed reads. To mitigate the effect of sequencing errors that are known to affect the Ion Torrent sequencing platform, we used a 99% OTU clustering threshold with a minimum of five reads per cluster (Loman et al., 2012; Salipante et al., 2014). Substantiation of the DNA metabarcoding method is required before its implementation for market samples. Therefore, here we have performed metabarcoding of blended formations of BM and CA. Blended formulations comprising 100% BM (BS1) showed 100% reads for BM, 100% CA (BS2) showed 100% reads for CA, 75% BM +25% CA (BS3) showed 79.4% reads for BM and 20.6% reads for CA, 50% BM +50% CA (BS4) showed 48.4% reads for BM and 51.6% reads for CA, 25% BM + 75% CA (BS5) showed 23.9% reads for BM and 76.1% reads for CA (Fig. 3, 4). This data demonstrated the accuracy of our metabarcoding pipeline and its suitability for market samples.

**Table 1.**
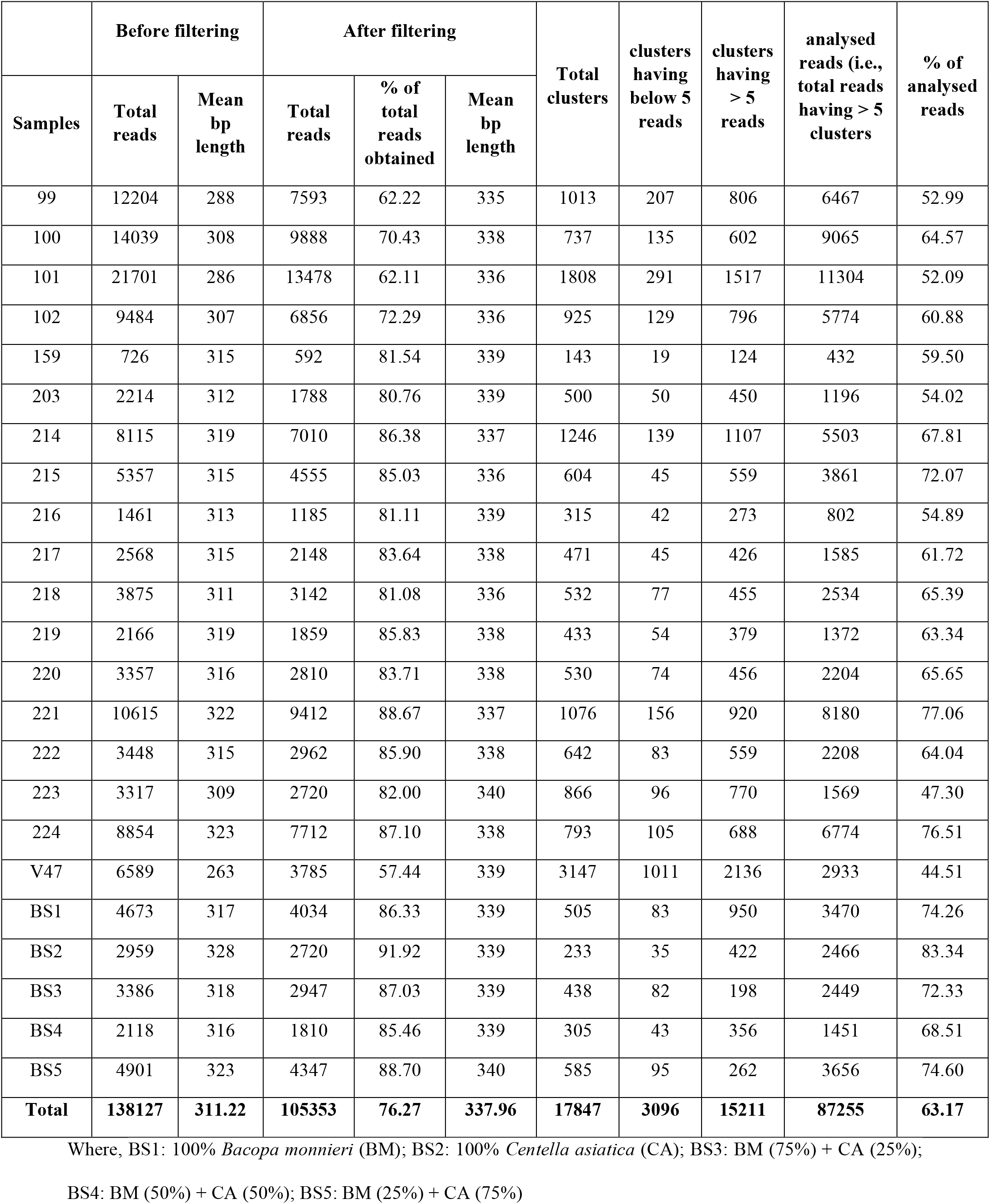
Raw data of DNA metabarcoding

**Figure 3.**
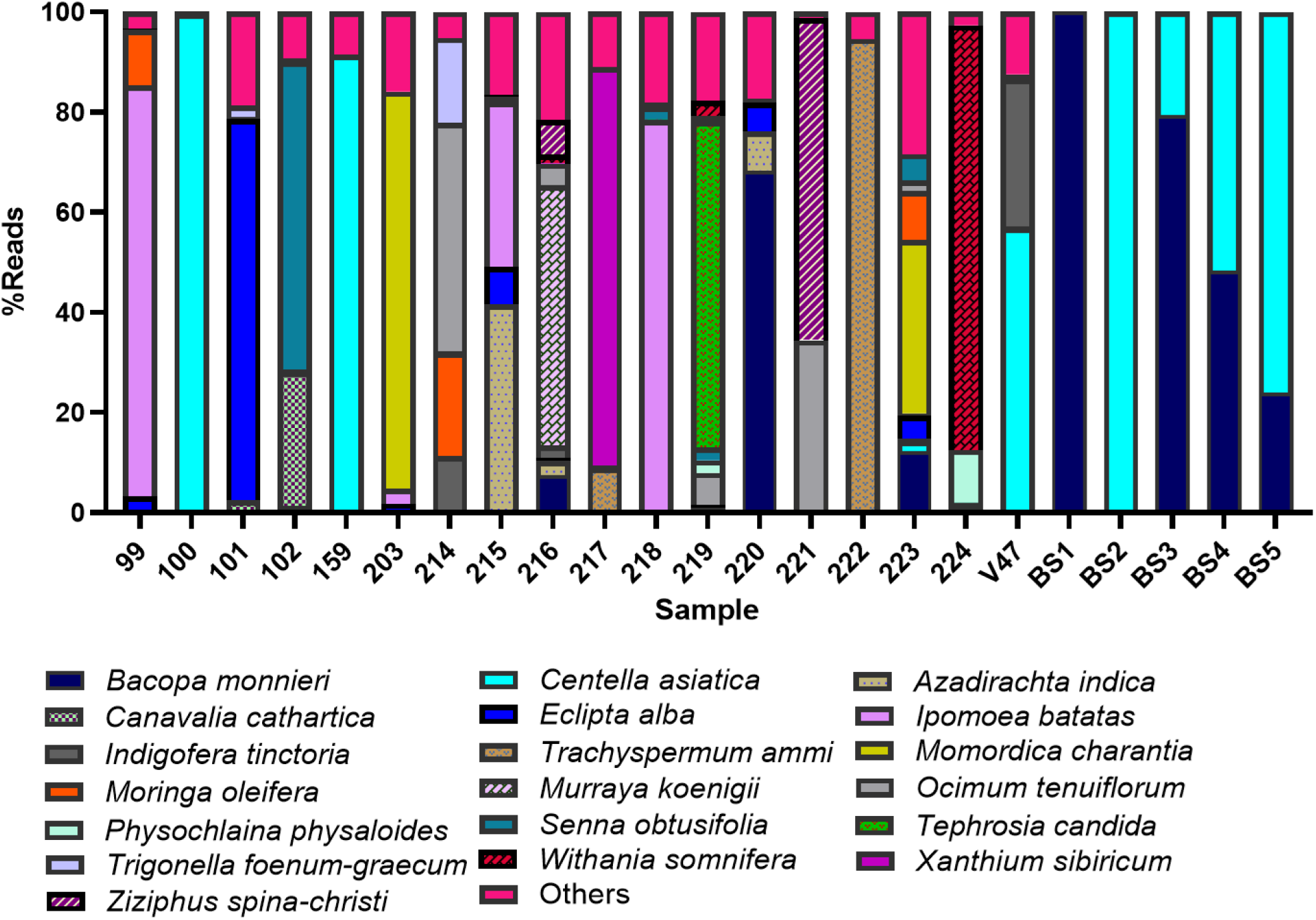
Relative abundance of botanicals at Genus level in Brahmi herbal products using DNA metabarcoding.

**Figure 4.**
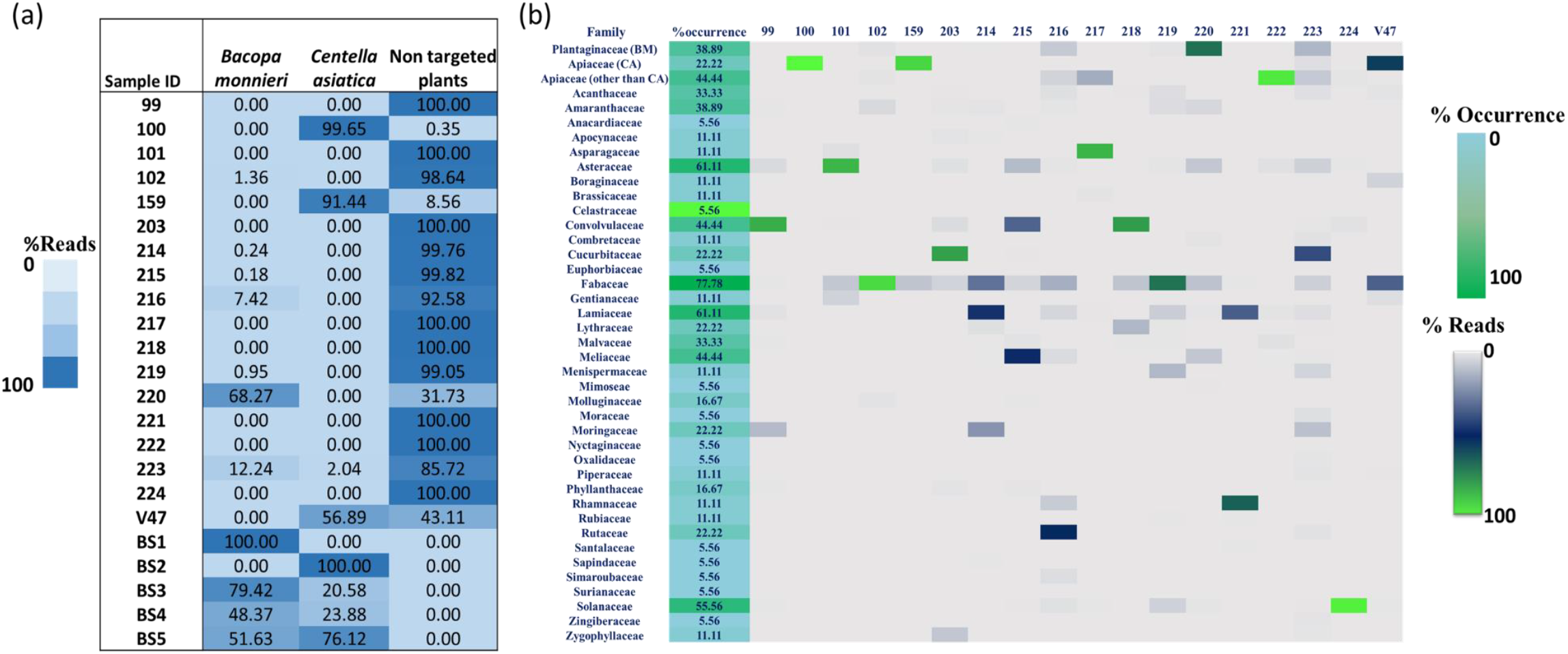
*Bacopa monnieri, Centella asiatica*, and other non-targeted botanicals detected in Brahmi herbal products (a) Heat map showing the relative abundance of targeted and nontarget botanicals present within Brahmi herbal products (b) Family level distribution of botanicals detected within Brahmi herbal products

A total of 119 species, 113 genera and 41 families have been identified from the market samples. Only four of the 18 market samples, labelled 100, 159, 223, and V47 showed 99.65%, 91.44%, 2.04% and 56.89% reads for CA, respectively, and five market samples labelled 102, 214, 215, 216, 219, 220, 223 showed 1.36%, 0.24%, 0.18%, 7.42%, 0.95%, 68.28% reads respectively for BM (Fig. 3, Fig. 4). A total of 12 different plant species were shown to be a prominent adulterant (covered >35% reads) in Brahmi market samples. For instance, sample 99 and 218 comprised 82.14% and 78.3% reads respectively for *Ipomoea batatas* (Family: Convolvulaceae); sample 101 comprised 76.12% reads of *Eclipta alba* (Family: Asteraceae); sample 102 comprised 61.6% reads of *Senna obtusifolia* (Family: Fabaceae); sample 203 and 223 comprise 79.3% and 34.6% reads respectively for *Momordica charantia* (Family: Cucurbitaceae); sample 214 comprised 45.6% reads of *Ocimum tenuiflorum* (Family: Lamiaceae); sample 215 comprised 41.36% reads of *Azadirachta indica* (Family: Meliaceae); sample 216 comprised 51.95% reads of *Murraya koenigii* (Family: Rutaceae); sample 217 comprised 79.5% reads of *Xanthium sibiricum* (Family: Asteraceae); sample 219 comprised 65.23% reads of *Tephrosia candida* (Family: Fabaceae); sample 221 comprised 64.3% reads of *Ziziphus spina-christi* (Family: Rhamnaceae); sample 222 comprised 94.38% reads of *Trachyspermum ammi* (Family: Apiaceae); sample 224 comprised 84.7% reads of *Withania somnifera* (Family: Solanaceae) (Fig. 3). Plant species and their percentage reads that are allied with others in Fig. 3 are listed in Table S3. The number of species detected per sample ranged from 2 to 28, with an average of 12.8 species in market samples.

A total of 44.4% samples exhibited 100% reads for non-targeted plants, 27.78% samples showed 90 to 99.8% reads for non-targeted plants, and 55.6% samples showed reads in the range of 0.18 to 99.65% for Brahmi (either BM or CA or both) (Fig. 4a). At family level, Fabaceae, Asteraceae, Lamiaceae, Solanaceae are prominent adulterants with presence in minimum 10 samples with 77.8 %, 61.1%, 61.1% and 55.5% occurrences (Fig. 4b). On the whole, this result emphasizes that a larger number of plant species have been detected in market herbal products while we did not observe non-targeted plant species in-house prepared blended formulations. The presence of unlabelled species might be deceiving, as DNA metabarcoding is one of the highly sensitive methods that can detect even trace amounts of contamination. For instance, pollen contamination because of wind-pollinated species during the stage of cultivation and harvesting, during transport, storage, production, and packing other plant species contaminated by inadequately cleaned containers, conveyors, and other equipment (Newmaster et al., 2013; Ivanova et al., 2016; Liu et al., 2018). However, DNA metabarcoding could only be applied for quality assessment to check the presence of labelled species and/or other non-listed botanicals, substitutes, and fillers in herbal products. The final results are influenced by a number of variables, including DNA quality and quantity proportional to the plant species and its biasness; DNA that can be removed or degraded during processing; PCR amplification bias due to poor primer fit and compatibility degree variation with different species; library preparation; sequencing platform; metabarcoding data analysis parameters; molecular identification algorithm; and reference databases (Pawluczyk et al., 2015; Staats et al., 2016). NCBI GenBank databases comprise non-curated databases which might lead to incorrect identification to the reference sequences at lower taxonomic level. However, NCBI GenBank can be used for identification at higher taxonomic levels (Hinchliff and Smith, 2014).

### Detection of chemical markers of BM and CA in market samples

A chemical marker-based authentication of BM and CA has been well demonstrated in the Indian pharmacopoeia (Indian pharmacopoeia 2007, volume 3). In the present study, the most commonly identified chemical markers bacoside A and asiaticoside were examined in Brahmi herbal products and blended formulations for validating DNA-based methods. Bacoside A is composed of four different triglycosidic saponins: bacoside A3, bacopaside II, bacopasaponin C, and the jujubogenin isomer, and it is the key bioactive constituent or chemical marker responsible for the pharmacological effects of BM, while asiaticoside is the most abundant triterpene glycoside found in the CA and is responsible for the pharmacological activities of CA (Deepak et al., 2005; Shinomol et al., 2011).

In HPLC chromatogram, a reference standard bacoside A gave four major peaks corresponding to bacoside A3 (1), bacopaside II (2), jujubogenin, isomer of bacopasaponin C (3), and bacopasaponin C (4) between retention time 20 to 25 min and reference standard asiaticoside gave a peak at retention time 5.08 min (Fig. S4). HPLC chromatogram of blended formulations showed peaks in the asiaticoside and bacoside A region which corresponds to their composition. For intense, 100% BM exhibited HPLC fingerprint at bacoside A region, 100% CA exhibited HPLC fingerprint at asiaticoside region, and admixture of BM and CA exhibited HPLC fingerprints of both (Fig. S5). Samples 102, 214, 216, 219, 220, and 223 showed similar qualitative HPLC fingerprints in the bacoside A region demonstrating the presence of BM. Samples 100, 159, and V47 showed a peak of asiaticoside obtained at a retention time (Rt) of 5.08 min demonstrating the presence of CA. The remaining nine samples (i.e. 50% samples) did not exhibit peaks corresponding to bacoside A or asiaticoside (Fig. 5).

**Figure 5.**
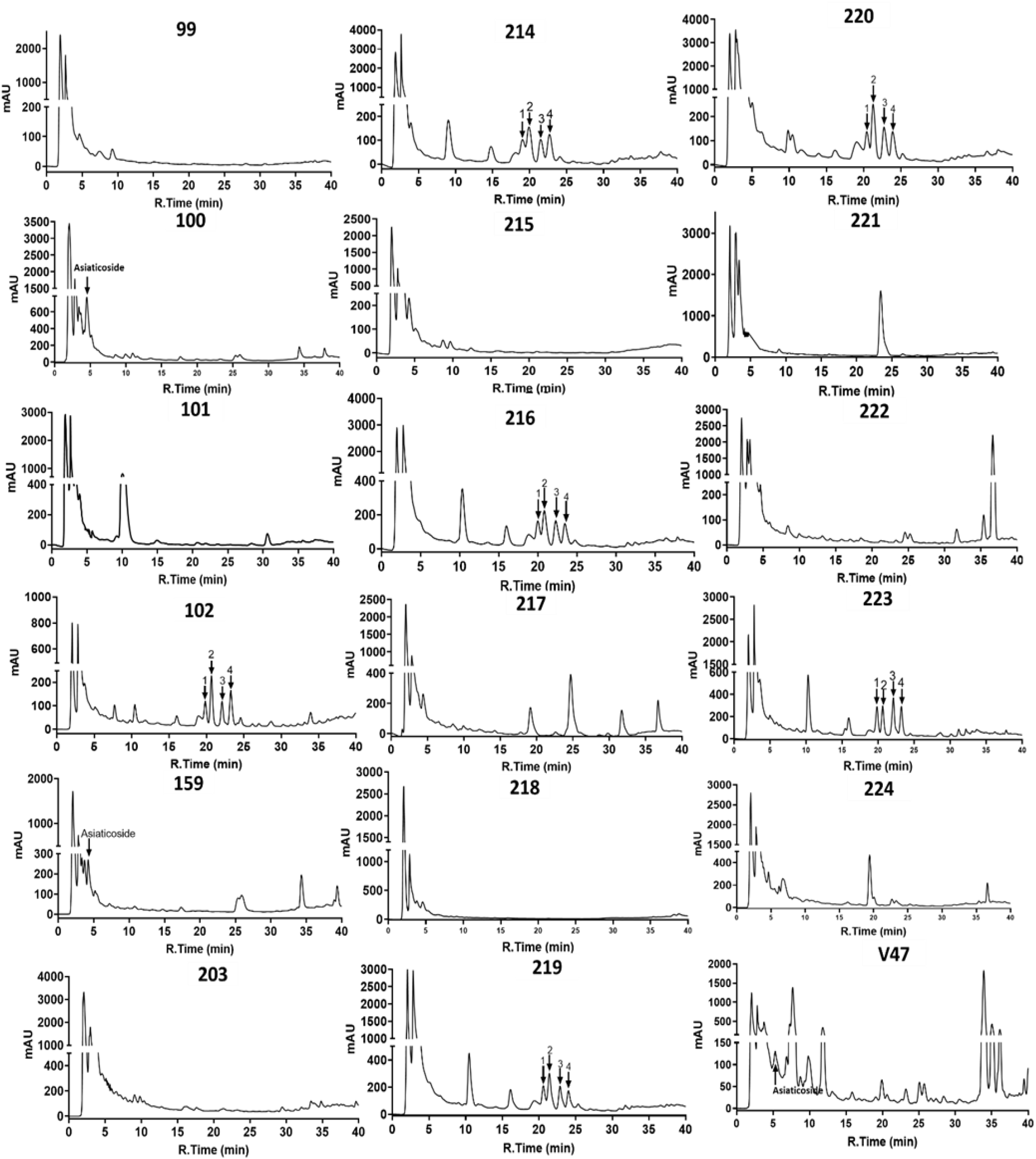
HPLC chromatograms of Brahmi herbal products. Bacoside A components Bacoside A3, Bacopaside II, Jujubogenin isomer of Bacopasaponine C, and Bacopasaponine C are indicated by arrows labeled 1, 2, 3, and 4 respectively. The peak of asiaticoside is marked with an arrow labeled with asiaticoside.

#### 3.1 Comparative results of all three approaches

Comparative results of PCR assay, DNA metabarcoding, and HPLC methods carried out in this study for detection of BM and CA in herbal products are represented in Table 2. In sample 223, BM and CA both are detected through metabarcoding, while by PCR-based approach and HPLC only BM was detected. In samples 102, 214, 216, and 219, BM was detected by HPLC and metabarcoding method but not through PCR-based approach and the presence of CA was detected by HPLC and metabarcoding method but not through PCR based approach in sample V47. This might be due to the availability of lower copy numbers of SCAR (used for BM detection)/ITS (used for CA detection) markers than *rbcL* markers, and in vati sample it could be due to the presence of impurities in the extracted DNA, as we observed lower *rbcL* gene amplification intensity in this sample (Fig. S1). Conclusively, presence of Brahmi (BM and CA) was detected in four samples (22.22%) by PCR based assay, in ten samples (55.56%) by metabarcoding and in nine samples (50.0%) by HPLC and in 8 samples (44.4%) BM and CA was not detected by all three methods. There were labels on market products for *Bacopa monnieri* on nine out of the 18 samples, but none had labels for *Centella asiatica* (Table S1). However only four of them had BM, while one of them had CA instead of BM. From the remaining 9 samples, where no species was mentioned on the label, BM and CA were found in 3 and 2 samples, respectively. The low level of fidelity raises concerns about the safety and reliability of Brahmi herbal products, especially in the context of discrepancies between labelling and constituents.

**Table 2.** Comparative result of PCR assay, DNA metabarcoding and HPLC

|  | PCR assay using BM and CA primers |  | Metabarcoding |  | HPLC |  |
| --- | --- | --- | --- | --- | --- | --- |
| Sample | *Bacopa monnieri* | *Centella asiatica* | *Bacopa monnieri* | *Centella asiatica* | Bacoside A (*Bacopa monnieri*) | Asiaticoside (*Centella asiatica*) |
| 99 | - | - |  | - | - | - |
| 100 | - | + | - | **+ (99.65% read)** | - | + |
| 101 | - | - | - | - | - | - |
| 102 | - | - | **+ (1.36% reads)** | - | + | - |
| 159 | - | + | - | **+ (91.44% reads)** | - | + |
| 203 | - | - | - | - | - | - |
| 214 | - | - | **+ (0.24% reads)** | - | + | - |
| 215 | - | - | **+ (0.18% reads)** | - | - | - |
| 216 | - | - | **+ (7.42% read)** | - | + | - |
| 217 | - | - | - | - | - | - |
| 218 | - | - | - | - | - | - |
| 219 | - | - | **+ (0.95% read)** | - | + | - |
| 220 | + | - | **+ (68.27% read)** | - | + | - |
| 221 | - | - | - | - | - | - |
| 222 | - | - | - | - | - | - |
| 223 | + | - | **+ (12.24% read)** | **+ (2.04 % read)** | + | - |
| 224 | - | - | - | - | - | - |
| V47 | - | - | - | **+(56.89% read)** | - | + |
| Absolute (relative) number of samples in which BM and CA was detected | 4.0 (22.22%) |  | 10.0 (55.56%) |  | 9.0 (50.0%) |  |

+ sign represents detection of respective plant species; - sign represents absence of respective plant species, % reads of BM and CA obtained in DNA metabarcoding is also represented.

## Conclusion

Comprehensive analysis of species-specific PCR, DNA metabarcoding, and HPLC for detection of BM and CA in herbal products revealed that DNA metabarcoding has benefits in terms of detection of a wide range of non-targeted adulterations. While HPLC is able to detect bioactive compounds, which are preferred in the identification and monitoring of the therapeutic potential of drugs, a straightforward species-specific PCR approach, which is rapid, affordable, and simple to analyse and handle, can overpower DNA metabarcoding and HPLC in terms of cost and experimental simplicity. From the collection of raw materials to the final furnished products, quality control of herbal products comprises various layers of investigation and authentication. Species-specific PCR, DNA metabarcoding, and HPLC approaches have distinct technological requirements, expenses, and levels of specialized knowledge required to provide a quality assessment of herbal products with defined applicability and inapplicability. The implementation of multiple or orthogonal authentication approaches, which are aligned to the concept of “fit-for-purpose,” is the need of the hour in quality control and safety of herbal products.

## Acknowledgment

Gujarat State Biotechnology Mission (GSBTM), Gandhinagar, Gujarat, India, has provided financial support for the project under the Research Support Scheme, grant ID GSBTM/JDRD/584/2018/204. The authors would like to thank Prof. Padamnabhi S. Nagar, The Maharaja Sayajirao University of Baroda, India for helping us in plant collection and authentication, Mr. Dipesh Parikh, Technical assistance, GBRC for carrying out HPLC, Mr. Nitin Savaliya, Technical assistance from Thermo Fisher Scientific, for NGS instrument handling and run setup.

## Author contribution

APS: Performed experiments, Data analysis, Writing and Editing manuscript, Validation of final manuscript; TT: Performed molecular biology experiments, Data analysis, Writing manuscript and validation; RP: Designed primers for metabarcoding, established metabarcoding data analysis pipeline, Manuscript editing; SS: Designed primers for CA, performed molecular biology experiments, Manuscript editing; CJ: Project administration, Methodology, Supervision, and Review & Editing; MJ: Principal Investigator, Conceptualization, Methodology, Supervision, and Review & Editing.

## Data availability statement

All data generated or analysed during this study are included in this article and its supplementary information files.

