## Supplementary material for "Comprehensive analysis using DNA metabarcoding, PCR, and HPLC unveils the adulteration in Brahmi herbal products": Figure S1, Table S1

### Supplementary materials

**Table S1.** Information about the herbal products

| Sample ID | Product name | Composition mention on labelled |
|---|---|---|
| 99 | Brahmi churna | Not mention |
| 100 | Brahmi powder | Not mention |
| 101 | Brahmi powder | Not mention |
| 102 | Brhami Churna | Not mention |
| 159 | Brhmi churna | *Bacopa monneri* |
| 203 | Brahmi powder | Not mention |
| 214 | Brahmi powder | Not mention |
| 215 | Brahmi powder | *Bacopa monnieri* |
| 216 | Brahmi leaf powder | Not mention |
| 217 | Brahmi powder | *Bacopa monnieri* |
| 218 | Brahmi leaf powder | *Bacopa monnieri* |
| 219 | Brahmi powder | *Bacopa monnieri* |
| 220 | Brahmi Churna | *Bacopa monnieri* |
| 221 | Brahmi leaf powder | *Bacopa monnieri* |
| 222 | Brahmi powder | *Bacopa monnieri* |
| 223 | Brahmi powder | *Bacopa monnieri* |
| 224 | Brahmi powder | Not mention |
| V47 | Brahmi Vati | Not mention |

**Table S2.** DNA concentration of the blended formations and herbal products

| Sample | DNA concentration (ng/µL) |
|---|---|
| 99 | 1.22 |
| 100 | 9.12 |
| 101 | 2.44 |
| 102 | 5.88 |
| 159 | 1.5 |
| 203 | 11.1 |
| 214 | 1.47 |
| 215 | 2.98 |
| 216 | 1.77 |
| 217 | 3.52 |
| 218 | 10.2 |
| 219 | 29.2 |
| 220 | 1.58 |
| 221 | 18.6 |
| 222 | 6.8 |
| 223 | 1.23 |
| 224 | 2.74 |
| V47 | 6.75 |
| BS1 | 5.5 |
| BS2 | 6.7 |
| BS3 | 4.9 |
| BS4 | 5.1 |
| BS5 | 4.7 |

Where, BS1: 100% *Bacopa monnieri* (BM); BS2: 100% *Centella asiatica* (CA); BS3: BM (75%) + CA (25%); BS4: BM (50%) + CA (50%); BS5: BM (25%) + CA (75%)

**Table S3.** Plant species and their % reads affiliated with others in Fig. 2

| Sample ID | Total % reads allied to others in Fig. 2 | Identified plants | Family | %reads |
|---|---|---|---|---|
| 99 | 3.400 | *Corchorus capsularis* | Malvaceae | 0.300 |
| | | *Cordia sebestena* | Boraginaceae | 0.200 |
| | | *Cyamopsis tetragonoloba* | Fabaceae | 0.100 |
| | | *Cyathula capitata* | Amaranthaceae | 0.500 |
| | | *Kinostemon ornatum* | Lamiaceae | 1.400 |
| | | *Phyllanthus emblica* | Phyllanthaceae | 0.400 |
| | | *Prosopis cineraria* | Fabaceae | 0.200 |
| | | *Pseudocaryopteris paniculata* | Lamiaceae | 0.100 |
| | | *Solanum lycopersicum* | Solanaceae | 0.200 |
| | | *Teucrium simplex* | Lamiaceae | 0.100 |
| | | *Vigna radiata* | Fabaceae | 0.100 |
| 100 | 0.353 | *Solanum nigrum* | Solanaceae | 0.121 |
| | | *Citrus polytrifolia* | Rutaceae | 0.099 |
| | | *Angelica sinensis* | Apiaceae | 0.066 |
| | | *Trigastrotheca stricta* | Molluginaceae | 0.066 |
| 101 | 18.825 | *Swertia cordata* | Gentianaceae | 5.263 |
| | | *Critonia sexangularis* | Asteraceae | 3.602 |
| | | *Biancaea sappan* | Fabaceae | 2.237 |
| | | *Asparagus racemosus* | Asparagaceae | 1.931 |
| | | *Conocliniopsis prasiifolia* | Asteraceae | 3.193 |
| | | *Guilandina bonduc* | Fabaceae | 0.984 |
| | | *Stenopadus talaumifolius* | Asteraceae | 0.687 |
| | | *Diplostephium alveolatum* | Asteraceae | 0.353 |
| | | *Libanothamnus occultus* | Asteraceae | 0.343 |
| | | *Vigna radiata* | Fabaceae | 0.121 |
| | | *Prosopis cineraria* | Fabaceae | 0.065 |
| | | *Gossypium sp.* | Malvaceae | 0.046 |
| 102 | 9.28 | *Alternanthera philoxeroides* | Amaranthaceae | 3.480 |
| | | *Angelica sinensis* | Apiaceae | 0.830 |
| | | *Biancaea sappan* | Fabaceae | 0.620 |
| | | *Glinus dahomensis* | Molluginaceae | 1.170 |
| | | *Lysiloma watsonii* | Fabaceae | 3.070 |
| | | *Neuontobotrys berningeri* | Brassicaceae | 0.100 |
| 159 | 8.565 | *Biancaea sappan* | Fabaceae | 8.560 |
| 203 | 15.95 | *Tribulus terrestris* | Zygophyllaceae | 8.210 |
| | | *Cyamopsis tetragonoloba* | Fabaceae | 4.000 |
| | | *Phyllanthus emblica* | Phyllanthaceae | 1.003 |
| | | *Gymnema sylvestre* | Apocynaceae | 1.000 |
| | | *Vigna radiata* | Fabaceae | 0.800 |
| | | *Oxybasis glauca* | Amaranthaceae | 0.512 |
| | | *Justicia adhatoda* | Acanthaceae | 0.420 |
| 214 | 5.233 | *Rotala rotundifolia* | Lythraceae | 1.981 |
| | | *Alternanthera philoxeroides* | Amaranthaceae | 0.981 |
| | | *Biancaea sappan* | Fabaceae | 0.636 |
| | | *Styphnolobium japonicum* | Fabaceae | 0.418 |
| | | *Rauvolfia serpentina* | Apocynaceae | 0.327 |
| | | *Sapindus mukorossi* | Sapindaceae | 0.273 |
| | | *Fabaceae sp.* | Fabaceae | 0.164 |
| | | *Distemonanthus benthamianus* | Fabaceae | 0.145 |
| | | *Boerhavia diffusa* | Nyctaginaceae | 0.127 |
| | | *Calliandra hygrophila* | Fabaceae | 0.091 |
| | | *Suriana maritima* | Surianaceae | 0.091 |
| 215 | 16.58 | *Aphanamixis polystachya* | Meliaceae | 6.010 |
| | | *Parthenium hysterophorus* | Asteraceae | 2.900 |

| | | | | |
|---|---|---|---|---|
| | | *Cyamopsis tetragonoloba* | Fabaceae | 2.850 |
| | | *Gossypium sp.* | Malvaceae | 1.090 |
| | | *Ambrosia trifida* | Asteraceae | 0.600 |
| | | *Phyllanthus emblica* | Phyllanthaceae | 0.600 |
| | | *Prosopis cineraria* | Fabaceae | 0.570 |
| | | *Cucumis melo* | Cucurbitaceae | 0.390 |
| | | *Sclerocarya birrea* | Anacardiaceae | 0.340 |
| | | *Trigastrotheca stricta* | Molluginaceae | 0.310 |
| | | *Vigna radiata* | Fabaceae | 0.260 |
| | | *Acalypha hispida* | Euphorbiaceae | 0.160 |
| | | *Corchorus olitorius* | Malvaceae | 0.130 |
| | | *Cressa cretica* | Convolvulaceae | 0.130 |
| | | *Ligusticum jeholense* | Apiaceae | 0.130 |
| | | *Medicago laciniata* | Fabaceae | 0.130 |
| 216 | 21.64 | *Falcataria moluccana* | Fabaceae | 9.8113 |
| | | *Angelica sinensis* | Apiaceae | 4.9057 |
| | | *Ailanthus altissima* | Simaroubaceae | 2.5157 |
| | | *Echinacanthus attenuatus* | Acanthaceae | 2.0126 |
| | | *Vigna radiata* | Fabaceae | 1.7610 |
| | | *Alternanthera philoxeroides* | Amaranthaceae | 0.6289 |
| 217 | 11.10 | *Trachyspermum ammi* | Apiaceae | 5.000 |
| | | *Ambrosia trifida* | Asteraceae | 4.400 |
| | | *Prangos trifida* | Apiaceae | 0.600 |
| | | *PREDICTED: Brassica* | Brassicaceae | 0.400 |
| | | *Malcolmia bicolor* | Brassicaceae | 0.400 |
| | | *Solanum nigrum* | Solanaceae | 0.300 |
| 218 | 18.114 | *Rotala rotundifolia* | Lythraceae | 12.076 |
| | | *Alysicarpus ovalifolius* | Fabaceae | 4.854 |
| | | *Desmodium styracifolium* | Fabaceae | 0.513 |
| | | *Justicia adhatoda* | Acanthaceae | 0.395 |
| | | *Sida szechuensis* | Malvaceae | 0.276 |
| 219 | 17.8 | *Tinospora sinensis* | Menispermaceae | 11.700 |
| | | *Alternanthera philoxeroides* | Amaranthaceae | 2.600 |
| | | *Andrographis paniculata* | Acanthaceae | 1.800 |
| | | *Echinacanthus attenuatus* | Acanthaceae | 0.500 |
| | | *Nesaea schinzii* | Lythraceae | 0.400 |
| | | *Kohautia caespitosa* | Rubiaceae | 0.400 |
| | | *Sida szechuensis* | Malvaceae | 0.400 |
| 220 | 17.44 | *Cicer arietinum* | Fabaceae | 6.447 |
| | | *Alternanthera philoxeroides* | Amaranthaceae | 3.803 |
| | | *Echinacea purpurea* | Asteraceae | 2.226 |
| | | *Urariopsis brevissima* | Leguminosae | 1.299 |
| | | *Glycyrrhiza aspera* | Fabaceae | 1.160 |
| | | *Terminalia chebula* | Combretaceae | 0.835 |
| | | *Solanum chacoense* | Solanaceae | 0.557 |
| | | *Leucaena retusa* | Mimoseae | 0.417 |
| | | *Aphanamixis polystachya* | Meliaceae | 0.371 |
| | | *Cajanus cajan* | Fabaceae | 0.325 |
| 221 | 1.300 | *Phylica nitida* | Rhamnaceae | 0.350 |
| | | *Jodina rhombifolia* | Santalaceae | 0.330 |
| | | *Amphidasya ambigua* | Rubiaceae | 0.320 |
| | | *Crotalaria podocarpa* | Fabaceae | 0.290 |
| 222 | 5.389 | *Conoclinium coelestinum* | Asterales | 1.585 |
| | | *Trachyspermum ammi* | Apiaceae | 1.495 |
| | | *Malva parviflora* | Malvaceae | 0.951 |
| | | *Sida szechuensis* | Malvaceae | 0.589 |
| | | *Platostoma chinense* | Lamiaceae | 0.408 |
| | | *Ambrosia trifida* | Lamiaceae | 0.362 |
| 223 | 28.43 | *Anethum graveolens* | Apiaceae | 7.393 |

| | | | | |
|---|---|---|---|---|
| | | *Luffa acutangula* | Cucurbitaceae | 4.207 |
| | | *Tinospora sinensis* | Menispermaceae | 2.422 |
| | | *Stephania tetrandra* | Menispermaceae | 2.358 |
| | | *Ficus amplocarpa* | Moraceae | 1.976 |
| | | *Citrus polytrifolia* | Rutaceae | 1.466 |
| | | *Andrographis paniculata* | Acanthaceae | 1.211 |
| | | *Sonchus asper* | Asteraceae | 1.020 |
| | | *Terminalia chebula* | Combretaceae | 1.406 |
| | | *Piper nigrum* | Piperaceae | 0.892 |
| | | *Alpinia pumila* | Zingiberaceae | 0.637 |
| | | *Strobilanthes cusia* | Acanthaceae | 0.637 |
| | | *Oxalis dillenii* | Oxalidaceae | 0.510 |
| | | *Tribulus terrestris* | Zygophyllaceae | 0.510 |
| | | *Glycyrrhiza aspera* | Fabaceae | 0.382 |
| | | *Rotala rotundifolia* | Lythraceae | 0.382 |
| | | *Solanum chacoense* | Solanaceae | 0.382 |
| | | *Desmodium renifolium* | Fabaceae | 0.319 |
| | | *Zingiber zerumbet* | Zingiberaceae | 0.319 |
| 224 | 2.75 | *Physalis chenopodifolia* | Solanaceae | 2.320 |
| | | *Conoclinium coelestinum* | Asteraceae | 0.310 |
| | | *Hansenia forbesii* | Apiaceae | 0.120 |
| V47 | 12.491 | *Borago officinalis* | Boraginaceae | 4.879 |
| | | *Swertia cordata* | Gentianaceae | 2.215 |
| | | *Styphnolobium japonicum* | Fabaceae | 2.111 |
| | | *Erythrostemon calycinus* | Fabaceae | 1.003 |
| | | *Justicia adhatoda* | Acanthaceae | 0.934 |
| | | *Moullava spicata* | Fabaceae | 0.623 |
| | | *Piper nigrum* | Piperaceae | 0.381 |
| | | *Lithospermum erythrorhizon* | Boraginaceae | 0.173 |
| | | *Monimopetalum chinense* | Celastraceae | 0.173 |

**Figure S1**

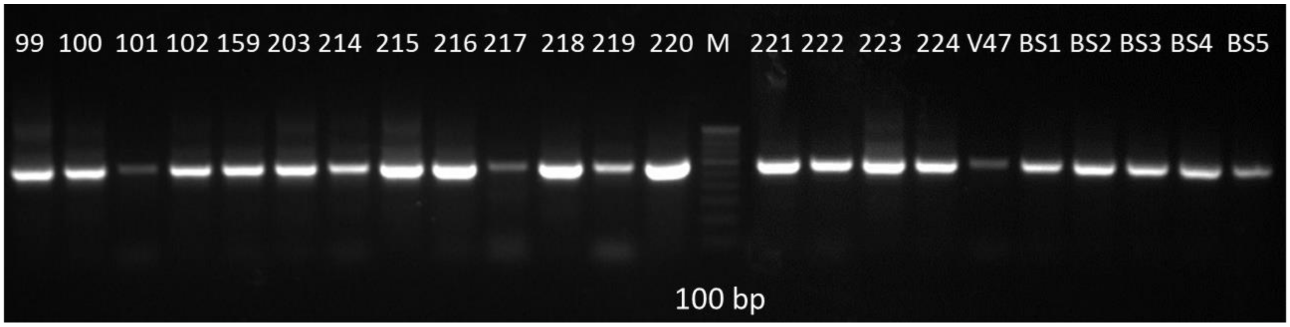


**Figure S1.** *rbcL* gene amplification of blended formations (BS1= 100% *Bacopa monnieri* (BM); BS2= 100% *Centella asiatica* (CA); BS3= BM (75%) + CA (25%); BS4= BM (50%) + CA (50%); BS5= BM (25%) + CA (75%)) and Brahmi market samples (99 to 102, 159, 203, 214 to 224, V47). M= 100 bp ladder.

**Figure S2**

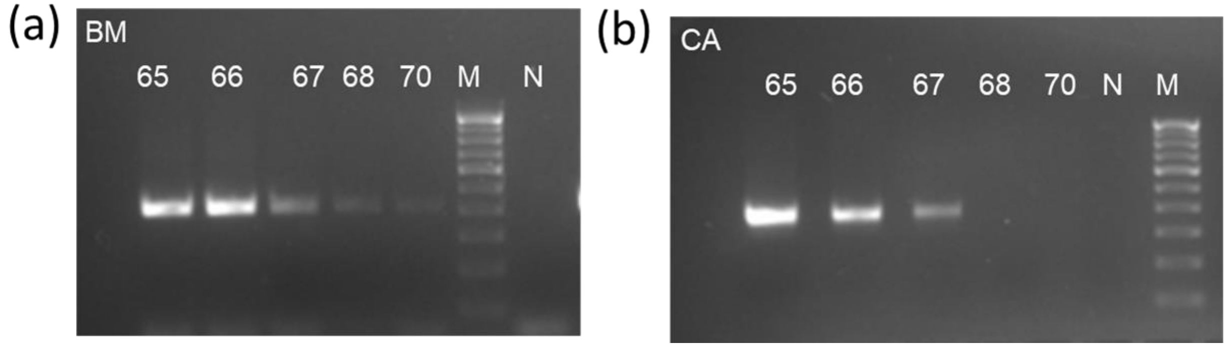


**Figure S2.** Optimization of PCR assay for *Bacopa monnieri* (BM) and *Centella asiatica* (CA) specific primers. Annealing temperatures 65, 66, 67, 68, and 70°C were used with respective plant DNA (10 ng). M=100 bp ladder, N= Non template control. (a) Optimization of annealing temperature for BM primers (b) Optimization of annealing temperature for CA primers.

**Figure S3**

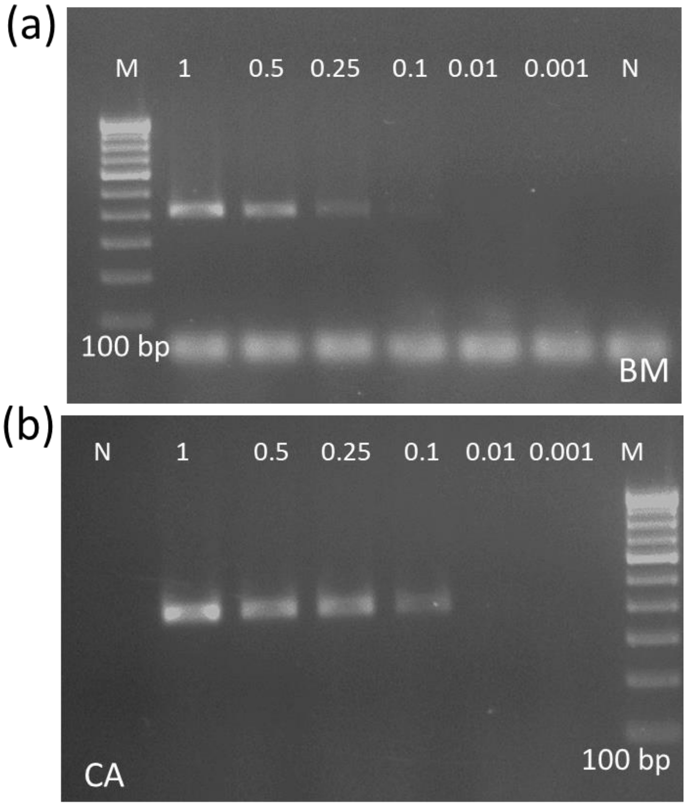


**Figure S3.** Sensitivity assay of the *Bacopa monnieri* (BM) and *Centella asiatica* (CA) specific primers. DNA concentrations 1.0, 0.5, 0.25, 0.1, 0.01 and 0.001 ng were used with respective primers in this assay. The amplicon size of both primers is 400 bp. M=100 bp ladder, N= Non template control (a) Sensitivity assay of the BM primers with BM plant DNA (b) Sensitivity assay of the CA primers with CA plant DNA.

#### Figure S4

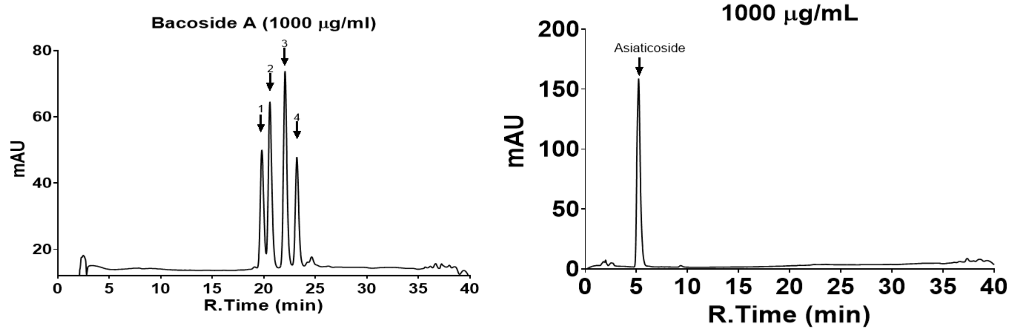


**Figure S4.** HPLC chromatograms of bacoside A and asiaticoside reference standard. 20 µL of 1000 µg/mL concentrations of both chemical markers were injected in HPLC. The chromatograms of bacoside A are on the left, and chromatograms of asiaticoside are on the right. Bacoside A components Bacoside A3, Bacopaside II, Jujubogenin isomer of Bacopasaponine C, and Bacopasaponine C are indicated by arrows labeled 1, 2, 3, and 4 respectively. The peak of asiaticoside is marked with an arrow labeled with asiaticoside.

**Figure S5**

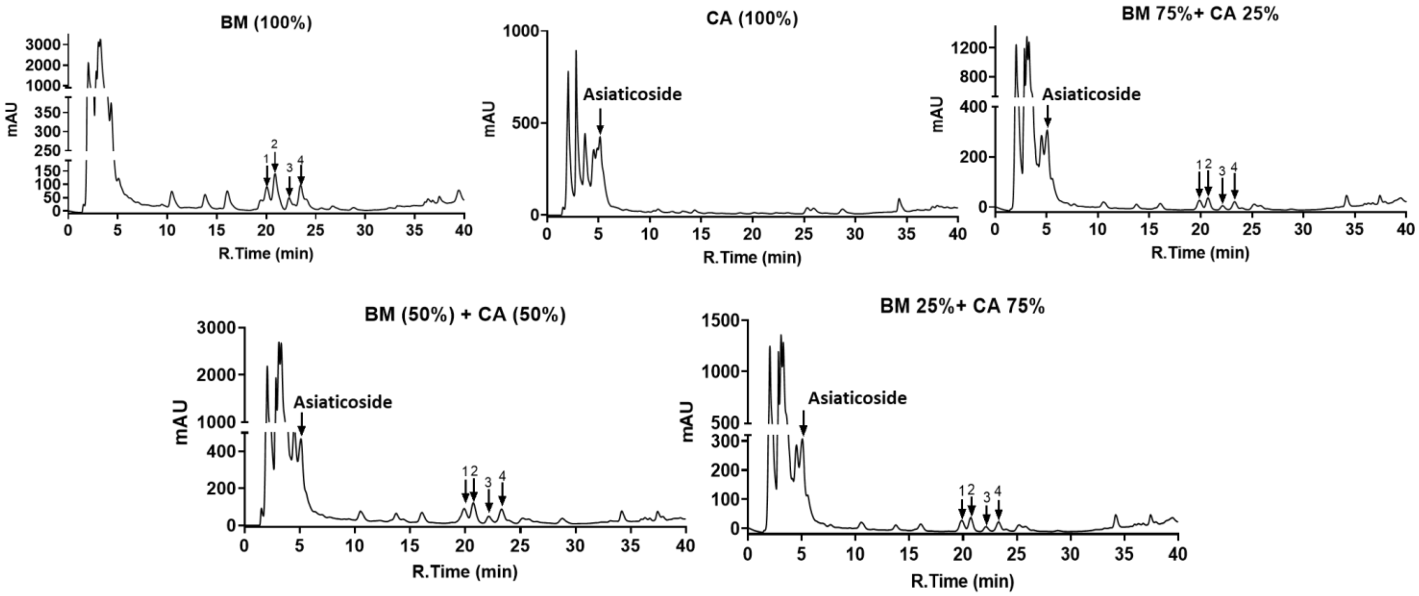


**Figure S5.** HPLC chromatograms of blended formulations. BM= *Bacopa monnieri* and CA= *Centella asiatica* (CA). Bacoside A components Bacoside A3, Bacopaside II, Jujubogenin isomer of Bacopasaponine C, and Bacopasaponine C are indicated by arrows labeled 1, 2, 3, and 4 respectively. The peak of asiaticoside is marked with an arrow labeled with asiaticoside.
